# First evidence towards chemical self-recognition in a gecko

**DOI:** 10.1101/2021.10.27.465717

**Authors:** Birgit Szabo, Eva Ringler

## Abstract

Self-recognition is the ability to recognise stimuli originating from oneself. Humans and most great apes show evidence of true self-recognition in the mirror test. They use their reflection to remove a mark that is only visible in the mirror. Not all animals, however, rely primarily on vision. In lizards, chemical cues are important in social interactions. A number of lizard species show chemical self-recognition but it has never been investigated in a gecko species. Here, we test the tokay gecko (*Gekko gecko*) a territorial species with parental care on their ability to discriminate their own skin and faecal chemicals from those of same-sex, unfamiliar conspecifics. Geckos show a higher response rate towards chemicals from unfamiliar individuals compared to self-produced chemicals and a water control. Lizards showed self-directed behaviour, responded stronger to skin chemicals and females responded more than males. Our study provides first evidence towards self-recognition and for a social function of chemical present on faeces in tokay geckos but further tests are needed to confirm true self-recognition. Tokay geckos are an excellent model species to investigate individual recognition to demonstrate more sophisticated social cognitive abilities than have previously been attributed to reptiles.

## Introduction

Self-recognition is the ability to recognise cues that represent/ originate from oneself (visual images, olfactory cues, acoustic stimuli) (Gallup, 1970; Gallup et al., 2011; Platek et al., 2004). Research into self-recognition aims to uncover self-awareness (the ability to become the object of one’s own attention) and its’ emergence across humans and non-human animals (Gallup, 1970; Gallup et al., 2011). The method of choice is the mirror self-recognition (MSR) test. In this test, a subject is confronted with a mirror and provided with a mark that can only be seen using the reflection in the mirror. Confirmation of MSR occurs when the subject inspects the mark and attempts to remove it using their reflection (Gallup, 1970). Two important control conditions need to be implemented. First, an invisible mark has to be used to exclude that physical irritation caused by the mark itself or the process of marking is triggering the behaviour (Gallup, 1970). Second, a mark has to be applied in a spot that can be seen without the use of the mirror to confirm the subjects’ motivation to remove marks in general (Gallup and Anderson, 2018). Humans and most great apes show MSR (Gallup, 1970; Gallup et al., 2011) while evidence in other species has led to controversial discussion (Gallup and Anderson, 2018; 2020).

Not all species primarily depend on their visual sense. This has led to the development of the sniff-test for self-recognition used in dogs whose primary sense is smell (Cazzolla Gatti, 2016). These studies have demonstrated that dogs discriminate between their own odour and that of conspecifics. They sniff the urine of unfamiliar dogs for longer than their own urine (Cazzolla Gatti, 2016; Horowitz, 2017). Furthermore, they sniff their own odour longer when it is modified than the modifier by itself (Horowitz, 2017). Nonetheless, some researchers have criticised these studies as not being a true equivalent to the MSR test because dogs do not show self-directed behaviour in the sniff-test which is an important criterion in the MSR test (Gallup and Anderson, 2018). Interestingly, chemical self-recognition tests are a fairly common test in lizards (Aguilar et al., 2009; Alberts, 1992; Aragón et al., 2001; Bull et al., 2000; Cooper, 1999; Graves and Halpern, 1991; López et al., 1997). Particularly one study in male desert iguanas (*Dipsosaurus dorsalis*) showed that these animals perform self-directed behaviour after detection of their own femoral gland sections but did not show this behaviour towards the secretions of unfamiliar males (Alberts, 1992). Following the critique voiced regarding the results in dogs, this study demonstrates more conclusive evidence for self-recognition using chemicals in lizards.

Generally, reptiles rely strongly on chemicals (i.e. pheromones) when it comes to individual recognition, territoriality, social interactions and mate choice (Norris and Lopez, 2001). In lizards, pheromones might originate from the skin or specialised glands such as femoral glands which are most pronounced in males (Norris and Lopez, 2001). Many species also possess cloacal glands that deposit pheromones onto the faeces (Norris and Lopez, 2001). This is especially important in scat piling lizards which defecate repeatedly in the same location (Bull et al., 1999a). Similar to latrines in mammals (e.g. Green et al., 2015; King et al., 2017), these scat piles can have a social function by communicating, for example, territory ownership (Bull et al., 1999a; 1999b) and group identity (Bull et al., 2000; but see Shah et al., 2006). Lizards detect pheromones using tongue-flicks (TF), protrusions of the tongue forward towards a stimulus (e.g. on the ground or on a swab) to collect chemicals (Cooper, 1994), and generally show increased TF rates towards stimuli from unfamiliar conspecifics (e.g. Alberts, 1992; Aragón et al., 2001; Cooper et al., 1999; Graves and Halpern, 1991).

Discrimination of self-produced chemicals and chemicals produced by unfamiliar, same-sex conspecifics has never been shown in a gecko species although leopard geckos (*Eublepharis macularius*) discriminate sex based on pheromones (Cooper and Steele, 1997; Mason and Gutzke, 1990) and thick-tailed geckos (*Nephrurus milii*) recognise their own scats to add additional faecal matter (Carpenter and Duvall, 1995). Many gecko species scat pile which suggests either a communicative function aimed at conspecifics, an anti-predatory function to avoid detection of refuges or both (Bull et al., 1999a; Carpenter and Duvall, 1995). Here, we test the tokay gecko (*Gekko gecko*), a large (up to 185 mm Snout Vent Length), nocturnal, insectivorous, scat piling gecko species from tropical South-East Asia (Grossmann, 2006). The aims of this study were to

1. investigate if tokay geckos can discriminate between self-produced chemicals and chemicals produced by unfamiliar, same-sex conspecifics on cotton swabs (Cooper, 1998).
2. We were also interested in finding out if chemicals originating from scats were as effective as chemicals originating from the skin as stimuli.

We predicted, that if geckos are able to recognize their own odour they would show lower responses towards their own odour than the conspecific odour (e.g. Alberts, 1992; Cooper et al., 1999; Graves and Halpern, 1991). If geckos are capable of self-recognition, we expected to find both stimulus directed and ground directed TFs as a sign of comparison between the two stimuli when confronted with their own and unfamiliar conspecific odour. We predicted, however, less ground directed responses when confronted with their own odour as it is familiar and can be recognised easier. We also predicted that faecal chemicals were as effective as skin chemicals in eliciting a response if scats had a communicative function.

## Methods

### Study animals, housing and husbandry

We tested 22 captive bred, adult tokay geckos, 10 males (SVL range = 11.35-15.02 cm) and 12 females (SVL range = 11.29-13.72 cm). Animals were acquired from different breeders across Europe and approximately 2-6 years old at the time of the study. Animals were naïve to the experimental procedure used in this study.

At our facility, geckos are kept singly in plastic terraria (females – 45 L x 45 B x 70 H cm; males – 90 L x 45 B x 100 H cm). Enclosures are equipped with a drainage layer of clay pebbles and a layer of organic rainforest soil (Dragon BIO-Ground) on top separated by a mosquito mesh to prevent mixing of the layers. On the soil surface we spread autoclaved red oak leaves. Collembola, isopods and earth worms in the soil break down the faecal matter produced by the geckos. Each enclosure also includes a compressed cork back wall, cork branches, refuges made out of cork branches cut in half and hung on the back wall as well as plants.

Enclosures are located in a fully controlled environment with a reversed photo period. Because tokay geckos are nocturnal, the dark cycle (when geckos are active) lasts from 6am to 6pm while the light period (when geckos are asleep) lasts from 6pm to 6am. Each enclosure is equipped with an additional light to provide lizards with UVB (Exo Terra Reptile UVB 100, 25 W) during the light cycle. The system automatically simulates a sunrise and sunset. Temperature is automatically control and reaches approximately 25 °C during the night cycle and 31 °C during the day cycle. To allow animals to thermoregulate, a heat mat (TropicShop) is fixed to the outside of each enclosure increasing the temperature by ∼5 °C. To simulate the tropical condition this species experiences in the wild, the room humidity is kept at 50% and daily rainfall (osmotic water, 30s every 12h at 5pm and 4am) increases the humidity within enclosures to 100%. Humidity decreases with time until the next rainfall event. All enclosures are set up on shelfs with small enclosures on the top and large enclosures on the bottom. Animals are spread evenly across two rooms.

Lizards are fed three times per week on Monday, Wednesday and Friday with adult crickets (*Acheta domesticus*). Before feeding, crickets are gut loaded using cricket mix (reptile planet LDT), Purina Beyond Nature’s Protein™ Adult dry cat food and fresh carrots to ensure that they provided optimal nutrition (Vitamin D and calcium). Each individual lizard receives 3-5 crickets each feeding with tweezers to be able to monitor the food intake. A water bowl provides water *ad libitum*. Once a month geckos are captured and weighed to ensure healthy weight.

### Experimental setup and stimuli

Lizards were tested in their home enclosures to reduce stress of handling (Langkilde and Shine, 2006) between 10^th^ of August to 30^th^ September 2021. Testing was conducted under red light (PHILIPS TL-D 36W/15 RED). The light we use has a red component at 718 nm which is not detectable by the tokay geckos’ photoreceptors (Loew 1994). Furthermore, a blue UV-C component at 282 nm is also produced which is visible to the geckos (Loew 1994) and promotes gecko activity (personal observation).

Because animals were spread across two rooms, each room was tested on a different, non-feeding day (either Tuesday or Thursday) once a week. Each individual was tested in a random order each day and the stimuli (control, own, same sex unfamiliar) and treatment (skin, faeces – i.e. scat) were also randomised across trials. As a positive control we used the odour of tap water on a paper towel. To create the control stimulus, one side of a cotton swab was taped 10 times on a moistened paper towel. As the familiar odour we used the individuals own odour either from their skin collected by gently rubbing one side of a cotton swab over its’ back and/or sides 10 times or from a fresh (no older than 2 days) scat. The cotton swab was rubbed on the scat until a stain was visible. To create the same-sex unfamiliar stimulus we took chemicals from the skin or scats of a same sex individual from the second room. Although animals never had direct contact with each other within a room we were unsure if the smell of individuals could spread within a room. To ensure true unfamiliarity, we used the individuals from the second room located across a small hallway. The same methods as for collecting individuals own odour was used. Each individual was tested on their reaction towards the odour of three different same-sex conspecifics. From each conspecific both chemicals from skin and faeces were used to be able to compare the reaction across treatments while controlling for identity. All cotton swabs were marked at the back to indicate on which side the stimulus was applied. This was done so the experimenter could present each cotton swab with the stimulus facing downwards to exclude the use of visual information originating from faeces or UV-reflecting chemicals (Mason 1992).

### Experimental procedure

At the start of a test day stimuli were set up as follows: First, all swabs for the control were prepared. Next, all swabs with lizards own odour were prepared and lastly, all swabs with the unfamiliar odour were prepared. This was done to leave enough time (20-30 minutes) between stimulus collection and test of focal individuals to recover from stimulus collection (skin treatment). All swabs were placed in clay bowls in the order of presentation (Figure 1). We first set up half of the individuals, tested their reaction and then set up and tested the second half. This was done to prevent excessive degradation of chemical stimuli. After all individuals finished testing, cotton swabs were discarded and clay bowls thoroughly cleaned with hot water and a sponge. Then they were dried upside down until the next test day. The experimenter ensured that the inside of the bowls was never touched. Furthermore, they ensured that the cotton swabs within a bowl never touched.

**Figure 1.**
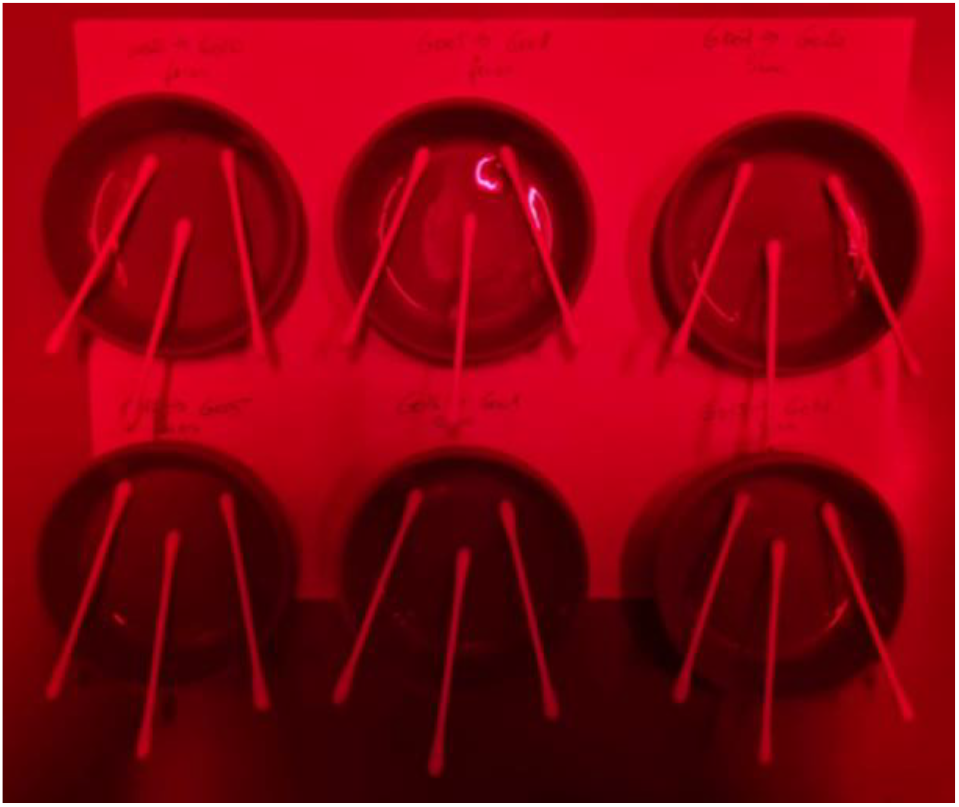
Setup of cotton swabs in clay bowls. For each focal subject swabs were placed in a separate clay bowl. Swabs were placed in the testing order. The experimenter made sure that swab tips covered in chemical stimuli never touched each other. To prevent excessive degradation of stimuli, individuals within a room were divided into two groups and the second group set up after the first group finished testing.

After set up, we first tested all individuals with the first cotton swab, then all with the second and finally with the third. This was done to leave about 10-15 minutes between stimulus presentations and reduce carry-over effects between stimuli. Both males and females were tested using the same procedure.

In a given trial, we first placed a dim white light (LED, SPYLUX LEDVANCE 3000K, 0.3 W, 17 lm) on top of the enclosure. Lizards were used to this light as it was used during feeding and generally during testing. Next, we located an individual in its’ enclosure. If the individual was hiding we gently removed the refuge from the back to expose it. Next, a cotton swab was presented holding it in a pair of 20 cm long metal tweezers. This ensured that the hand of the experimenter was far enough away to prevent the experimenters’ odour interfering with the experiment. The experimenter was visible during trials similar as during regular feeding. Trials from the first two test days were recorded with a GoPro (Hero 5). However, videos were too dark and we had issues scoring the lizards behaviour. For all other trials we switched to recording with a Samsung S20 smartphone (108 Megapixel, 8K-FUHD) which produced far better quality videos under the light conditions. Furthermore, the ability to switch between front and back camera enabled us to take videos even when lizards were sitting above the tank entrance. By the second week of testing we detected a large decrease in bites likely caused by lizards learning that the cotton swab was not edible. We, therefore, decided to repeat the first trial at the end of the testing period to ensure that our measurements were not confounded by changes in behaviour.

### Data collection

Videos were scored from the start of a trial, when the cotton swab was first presented within 1 cm of the lizards snout. Trials lasted a maximum of 120 seconds (2 minutes). If a tongue flick (TF) or bite occurred, the behaviour was video recorded for 60s after the initial event following the procedure used in previous studies with squamates (e.g. Aragón et al., 2001; López et al., 1997; Martin et al. 2020). If the lizard showed a turn (whole body movement away from the swab, Table 1) and walked away from the swab the trial was terminated.

**Table 1.** Ethogram of behaviours shown by tokay geckos in response to chemical stimuli.

| Name of behaviour | Description |
| --- | --- |
| Breath | One up and down movement of the lizards' throat = 1 breath. Only visible from the ventral side. Recorded as counts. |
| Deep breath | One extension and retraction of the flanks behind the front legs. Visible from the dorsal and ventral side. Recorded as counts. |
| Tongue flick | Quick protrusion of the tongue forward away from the mouth. NOT licking of the lips which is also a protrusion of the tongue but along the skin of the mouth. Recorded as counts. |
| Bite | The tip of the swab is taken between the upper and lower jaw. May be accompanied by shaking of the head. Recorded as counts. |
| Turn | The lizards' moves away from the swab. The whole body moved either past the swab, backwards away or involved a turn away from the swab. Recoded as yes or no. A trial was terminated if this behaviour was shown. |

Videos were analysed blind as to which stimulus was presented but not to treatment which was visible in some videos. We used VLC media player (Version 3.0.7.1, Vetinari, Intel 64 bit) to score behaviour (Table 1) shown during trials. We scored bites, TF, breaths if the ventral side of the individual was visible (i.e. gular pumping, Norris and Lopez, 2011), deep breaths, and turns. TFs were divided into flicks directed at the stimulus (tongue tip aimed at the swab) and flicks directed towards the ground (tip aimed at the ground). We also measured the trial time in seconds starting from the time the stimulus was presented within 1 cm of the lizards’ snout until either 120 seconds without a bite or TF elapsed, the lizard performed a turn or 60 seconds after the first bite or TF. Enclosure temperature was recoded automatically to an accuracy of 0.1 °C by the system responsible for regulating the environment within rooms.

### Ethical note

The test reported in this study are strictly non-invasive observations of behaviour. Introducing odour of conspecifics is a practice used during enrichment in reptiles. We followed the guidelines provided by the Association for the Study of Animal Behaviour/ Animal Behaviour Society for the treatment of animals in behavioural research and Teaching (2018). All testing was approved by the Suisse Federal Food Safety and Veterinary Office (National No. 33232, Cantonal No. BE144/2020). Captive conditions were approved by the Suisse Federal Food Safety and Veterinary Office (Laboratory animal husbandry license: No. BE4/11).

### Statistical analysis

#### Power analysis

Before data collection, we performed a power analysis using G^*^power (Faul et al., 2007; 2009). As our study was designed as a 2×2×3 factorial designed, we calculated power based on a within factor repeated measures ANOVA. The literature on chemical discrimination in other lizard and worm lizard species (Alberts,1992; Cooper et al., 1999; López et al., 1997) generally suggested large effect sizes. We were, however, unsure what effect size to expect from our geckos and therefore calculated the minimal effect size that could be reliably detected at a power of 0.8. We specified an alpha level of 0.05, a power of 0.8, six groups with three measurements, a correlation among repeated measures of 0.5 and a correction of 1. With these settings and a sample size of 24 individuals we are able to detect an effect size of 0.3 at an actual power of 0.99. The sample size used in our study was 22 individuals. We expected, however, only a slight reduction in the actual power to detect a small effect size.

#### Data analysis

### General reaction

We were interested if the reaction towards the presented stimuli was affected by the treatment (scat or skin), stimulus (water control, own or unfamiliar odour), sex (male or female), the order in which stimuli were presented, trial and temperature. These were used as fixed effects in three models looking at all tongue flicks produced in 60 seconds, deep breaths per second and breaths per second. For TFs we used a generalised linear mixed zero-inflation Poisson model (GLMM, package glmmTMB, Brooks et al., 2017) because our dataset included a large amount of 0 TFs. The conditional model included the above mentioned fixed effects while the zero-inflation model only included treatment, stimulus and sex as fixed effects. We did not expect all fixed effects to cause zero-inflation. Both the models included a random effect of animal identity to account for repeated measures. We were also interested if the difference in size between the test subject and the unfamiliar individual (delta SVL) from which the odour was taken affected TFs. To this end, we looked at TFs produced in the unfamiliar condition only as the response variable in another zero-inflation Poisson model. Both the conditional model and zero-inflation model included delta SVL and treatment as fixed effects and trial and animal identity as random effects. In both analyses we specified session for the over-dispersion component.

For the two measures of breathing, we first divided the number of breaths by the trial time to get a comparable measure for the breaths (breaths and deep breaths per second). We used breaths and deep breaths per second as the response variable in linear mixed effects models (LME, package lmerTest, Kuznetsova et al., 2017) with Gaussian family including the above mentioned fixed effects. Both models conformed to the assumption of residual normality (visual inspection of qqplots). Both models included a random effect of animal identity and session to account for repeated measures. We did not analyse bites because they were shown too infrequent to be analysed.

### Differences between swab and ground directed tongue flicks

Across all trials, males only tongue flicked three times while females together produced 202 TFs. We, therefore, based the following analysis on the data from females only. To identify if lizards compared their own odour to that of an unfamiliar individual we recorded TFs directed at the swab and those directed at the ground (on which their own odour was present). We analysed swab and ground directed TFs separately. We used generalised linear mixed zero-inflation Poisson models with ground or swab TFs as the response variable. The conditional models included stimulus as the only fixed effect as well as treatment, trial and animal identity as random effects. Treatment and trial were included as random effects because the general analysis revealed significant effects on TFs. In the zero-inflation models we used treatment and stimulus as fixed effects and animal identity as the random effect. We also specified session for the over-dispersion component. Finally, we also compared the two TFs within stimulus conditions using their average across trials and treatments (to avoid pseudo-replication) with Wilcoxon signed rank tests for paired data.

Data analysis was done in the free, open source software R (Version 4.0.3; R Core Team, 2020). All data and code produced during this study are available on the Open Science Framework (OSF; link for review purposes: https://osf.io/jp7h8/?view_only=b4c0eac3792f4adaaef2ff6745aebf45)

## Results

One female (G015) could not be tested as she was too anxious and was only used as a stimulus individual. All other geckos habituated fast to being rubbed on their back with a swab and did not flee during stimulus collection by the second week of testing (the first week of testing was not used for analysis).

### General reaction to the presented stimuli

Our analysis revealed that males tongue flicked much less than females (GLMM, estimate = - 4.249, *z*-value = -4.674, CI_low_ = -6.031, CI_up_ = -2.467, *p*-value < 0.001; Figure 2A) and that lizards tongue flicked less towards odour originating from scats than from skin (GLMM, estimate = -0.406, *z*-value = -2.413, CI_low_ = -0.736, CI_up_ = -0.076, *p*-value = 0.016, Figure 2B). Compared to swabs containing the odour of unfamiliar individuals, lizards directed less TFs towards tap water from a paper towel (GLMM, estimate = -0.556, *z*-value = -2.733, CI_low_ = - 0.954, CI_up_ = -0.157, *p*-value = 0.006; Figure 2A) and their own odour (GLMM, estimate = - 0.698, *z*-value = -3.562, CI_low_ = -1.083, CI_up_ = -0.314, *p*-value = 0.0004; Figure 2A). Overall, lizards decreased TFs over the course of the experiment (GLMM, estimate = -0.196, *z*-value = -2.099, CI_low_ = -0.378, CI_up_ = -0.013, *p*-value = 0.036). We detected no effect of stimulus order (GLMM, estimate = 0.043, *z*-value = 0.481, CI_low_ = -0.133, CI_up_ = 0.219, *p*-value = 0.631) and temperature (GLMM, estimate = 0.086, *z*-value = 0.277, CI_low_ = -0.524, CI_up_ = 0.697, *p*-value = 0.782) and the size of the stimulus individual had no effect on the number of TFs (GLMM, estimate = -0.123, *z*-value = -0.667, CI_low_ = -0.524, CI_up_ = 0.697, *p*-value = 0.505). The zero-inflation models did not produce any significant results (GLMM, *p*-value > 0.05; Table A1 and A2).

**Figure 2.**
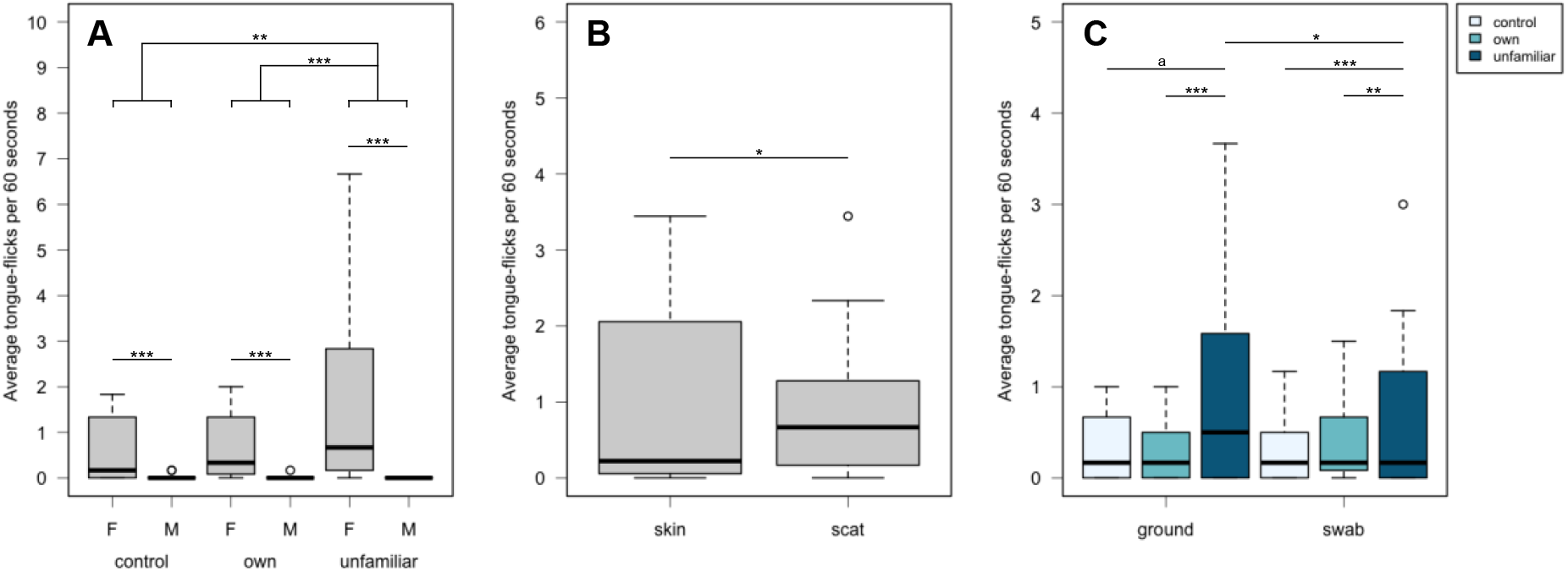
(A) Average tongue flicks produced within 60 seconds in the three stimulus conditions separated between males and females. Overall, lizards tongue flicked the most in the unfamiliar conditions and males tongue flicks less than females. (B) Average tongue flicks produced within 60 seconds across the skin and scat treatment. Lizards tongue flicked less in response to scats. (C) Average tongue flicks produced within 60 seconds directed towards the ground and the swab separated into stimulus conditions. Only data from females are shown. Females tongue flicked more when stimuli originated from unfamiliar individuals (dark blue) compared to the own (medium blue) and control condition (light blue). Females tongue flicked the ground more than the swab in the unfamiliar condition only. a *p* < 0.06, ^*^ *p* < 0.05, ^**^ *p* < 0.01, ^***^ *p* < 0.001.

We found no significant effects of any of the fixed effects on deep breaths per second (LME, *p*-value > 0.05, Table A3) but geckos showed a higher breathing rate in response to stimuli originating from scats (when responses towards all stimuli were lumped together) indicating a stronger involvement of olfaction (LME, estimate = 0.106, *t*-value = 3.132, CI_low_ = 0.039, CI_up_ = 0.170, *p*-value = 0.002, Table A4). None of the other fixed effects were significant (LME, *p*-value > 0.05, Table A4).

### Differences between swab and ground directed tongue flicks

Females directed less ground TFs towards their own odour compared to odour from unfamiliar individuals (GLMM, estimate = -1.133, *z*-value = -3.872, CI_low_ = -1.707, CI_up_ = -0.560, *p*-value = 0.00011, Figure 2C). Females also directed less swab TFs towards their own odour (GLMM, estimate = -0.927, *z*-value = -2.979, CI_low_ = -1.538, CI_up_ = -0.317, *p*-value = 0.003, Figure 2C) and even less towards the water control (GLMM, estimate = -1.405, *z*-value = -4.614, CI_low_ = -2.002, CI_up_ = -0.808, *p*-value < 0.001, Figure 2C). The zero-inflation models did not produce any significant results (GLMM, *p*-value > 0.05, Table A5 and A6). Within conditions females directed more TFs towards the ground in the unfamiliar condition (Wilcoxon signed rank tests, V = 3, N = 11, *p*-value = 0.012, Figure 2C). We found no differences in the control (Wilcoxon signed rank tests, V = 18.5, N = 11, *p*-value = 0.801, Figure 2C) and own odour condition (Wilcoxon signed rank tests, V = 14.5, N = 11, *p*-value = 0.829, Figure 2C).

## Discussion

Our experiment demonstrated that, at least female tokay geckos, discriminate between their own odour and that of an unfamiliar female using chemicals originating from the skin and scats but they show a weaker response to chemicals originating from scats. More TFs occurred towards the odour of unfamiliar individuals and females produced more ground directed TFs in response to the unfamiliar conspecific odour. In general, lizards sampled the stimulus most (swab directed TFs) in the unfamiliar condition and we found no differences in TFs directed towards their own odour and the water control.

Based on previous studies in other lizards (e.g. Alberts, 1992; Cooper et al., 1999; Graves and Halpern, 1991), we predicted that tokay geckos would show more TFs towards chemical stimuli originating from unfamiliar, same-sex conspecifics. Our results are in line with these studies but only in females. Males only tongue flicked a total of three times during the course of the experiment. Either, males do not rely as strongly on skin and scat chemicals for individual recognition or they show a delayed response which we did not record using our methodology. Indeed, we observed an increase in activity including TFs in some males after trials had ended. Male tokay geckos are territorial (Grossmann, 2006) and their behaviour might be interpreted as searching for the intruder. It is, however, necessary to run additional test recording not just the immediate response of males within two minutes but record behaviour for a longer time such as 10-15 minutes after stimulus presentation. Furthermore, males might react stronger to femoral gland secretions similar to male amphisbaenians (*Blanus cinereus*; Cooper et al., 1994) which should be tested in the future.

We also predicted that lizards would show chemical self-recognition by producing more ground directed TFs in response to the unfamiliar odour compared to their own odour. Our results confirm our prediction. Testing individuals inside their own enclosure posed an experimental advantage. Their enclosures are saturated with their own odour which made it possible to detect “self-directed” behaviour which would not have been possible in a neutral environment. Although male desert iguanas showed self-directed TFs towards their femoral glands (Alberts, 1992), we did not expect to find such behaviour in our geckos. Tokay geckos are, however, territorial, show site fidelity and scat pile. We expected, therefore, that if any comparison between the presented stimuli and self-produced odour did take place, this would likely be shown by TFs towards the ground. These ground directed TFs were very pronounced and easy to score because animals would always turn their heads away from the swab before tongue flicking the ground. Our results point towards similar difficulty to recognise tap water and their own scent because ground and swab directed TFs did not differ across these test conditions. They did differ in the unfamiliar condition. We recorded higher rates of ground directed TFs compared to swab directed TFs demonstrating a need for increased comparison with their own odour. Interestingly, a study in male Iberian rock-lizards (*Lacerta monticola*) showed no differences in non-swab directed TF between males own and unfamiliar males femoral gland secretion (Aragón et al., 2001). This study tested wild caught males that were kept together with a second individual on their reaction to femoral gland secretions. We used chemical from skin and scats from captive bred individuals kept singly and mainly analysed the reaction from females to these stimuli. It is possible that the scent of the second individual present in the enclosure interfered with “self-directed” TFs in rock-lizards. A comparison to our results is, however, difficult due to the many differences between studies.

A next step towards more conclusive evidence for true self-recognition would be to test geckos’ reaction towards a change in their own odour similar to what was done with dogs (Horowitz, 2017). Dogs are more interested in their own odour when it was marked but where less interested in the mark alone. If geckos similarly increase ground directed TFs towards their marked odour compared to the mark alone then this would further support our geckos’ ability to show true self-recognition.

Previous studies have considered that an increased rate of TFs towards the odour of unfamiliar individuals could be caused by novelty of the stimulus (Bull et al., 1999a; 2000). Bull and colleagues (1999a; 2000) used chemical stimuli from heterospecific individuals that were unfamiliar to the test lizards as a control. If novelty was causing increased TF rates then lizards would also show an increased response towards the heteropsecifics’ odour which they did not ruling out novelty as a cause for increased TF rates (Bull et al., 1999a; 2000). In our study, we used tap water from a paper towel as a control instead of odourless deionised water which elicited a lower rate of TFs compared to the unfamiliar odour. The fact that similar numbers of TFs (ground and swab directed) were directed towards their own odour and the odour of tap water and a paper towel shows that novelty was not solely responsible for our lizards’ reaction. As tap water and paper towels are not odourless, we would expect increased TF rates to inspect the novel odour which we did not find. We acknowledge, however, that an additional control similar to what was used in previous studies (Bull et al., 1999a; 2000) is needed to completely rule our novelty as a cause for the strong effect we found. Furthermore, we can also rule out that diet differences caused the difference in response towards own and unfamiliar odour because all our lizards were fed the same diet.

Finally, our results also point towards a social function of scat piling. Although geckos produced less TFs towards scats this difference was small. Additional research could determine if geckos inspect scat piles of other individuals when available, if they are more likely to defecate in locations with their own scat present (Carpenter and Duvall, 1995), and could investigate if lipids are deposited on scats by glands (Bull et al., 1999b). Furthermore, scat piling might have a possible function related to predator avoidance when predators use the odour of scats to locate refuges (Bull et al., 1999a; Carpenter and Duvall, 1995; Norris and Lopez, 2011). Studies on wild lizards should document the location of scat piles to determine if scat piles have an anti-predator function as well. Scat piles in locations that are not frequently visited by geckos would point towards an anti-predator function.

In summary, we provide fist evidence for chemical self-recognition in a gecko species and a possible social function of scat piles. Further investigations are, however, needed to confirm true self-recognition in tokay geckos and to better understand the communicative function of scats. Future studies could also look at other forms of recognition such as discrimination between familiar and unfamiliar individuals, mate recognition and kin recognition of skin, femoral gland and scat odours. Tokay geckos are a good model species to investigate recognition in general as they show biparental care and form temporary family groups with their offspring (Grossmann, 2006; Somma, 2003). Such studies can potentially demonstrate more sophisticated social cognitive abilities than have previously been attributed to reptiles (Doody et al., 2013).

## Acknowledgements

This work was supported by the University of Bern, the Austrian Science Fund (FWF) [grant P 31518, PI: ER] and the Swiss National Science Foundation (SNSF) [grant 197921, PI: ER]. We would like to thank Eva Barbara Zwygart for her support with animal husbandry.

## Appendix Results tables

**Table A1.** Parameter estimates and test statistics for the generalised linear mixed zero-inflation Poisson model looking at all tongue flicks produced by all tested individuals. The models included a random effect of animal identity and an over-dispersion parameter of session. Significant results are highlighted in bold. CI – confidence interval.

| Conditional model |  |  |  |  |  |
| --- | --- | --- | --- | --- | --- |
| Parameter | Estimate | z-value | CI<sub>low</sub> | CI<sub>up</sub> | p-value |
| Intercept | -1.924 | -0.239 | -17.681 | 13.833 | 0.811 |
| **Scat** | **-0.406** | **-2.413** | **-0.736** | **-0.076** | **0.016** |
| **Water control** | **-0.556** | **-2.733** | **-0.954** | **-0.157** | **0.006** |
| **Own odour** | **-0.698** | **-3.562** | **-1.083** | **-0.314** | **0.0004** |
| **Male** | **-4.249** | **-4.674** | **-6.031** | **-2.467** | **2.95\*10<sup>-6</sup>** |
| Stimulus order | 0.043 | 0.481 | -0.133 | 0.219 | 0.631 |
| **Trial** | **-0.196** | **-2.099** | **-0.378** | **-0.013** | **0.036** |
| Temperature | 0.086 | 0.277 | -0.524 | 0.697 | 0.782 |
| Zero-inflation model |  |  |  |  |  |
| Parameter | Estimate | z-value | CI<sub>low</sub> | CI<sub>up</sub> | p-value |
| Intercept | -1.462 | -1.083 | -4.107 | 1.184 | 0.279 |
| Scat | -1.959 | -1.820 | -4.069 | 0.151 | 0.069 |
| Water control | 1.393 | 1.282 | -0.737 | 3.523 | 0.200 |
| Own odour | 0.469 | 0.450 | -1.574 | 2.511 | 0.653 |
| Male | -17.667 | -0.003 | 0.554 | 5.785 | 0.998 |

**Table A2.** Parameter estimates and test statistics for the generalised linear mixed zero-inflation Poisson model looking at all tongue flicks produced in the unfamiliar condition by all tested individuals. The models included trial and animal identity as random effects. Significant results are highlighted in bold. CI – confidence interval.

| **Conditional model** |  |  |  |  |  |
| --- | --- | --- | --- | --- | --- |
| **Parameter** | **Estimate** | **z-value** | **CI<sub>low</sub>** | **CI<sub>up</sub>** | **p-value** |
| **Intercept** | **-2.662** | **-2.233** | **-4.998** | **-0.325** | **0.026** |
| Scat | -0.318 | -1.405 | -0.761 | 0.126 | 0.160 |
| Delta SVL | -0.123 | -0.667 | -0.485 | 0.239 | 0.505 |
| **Zero-inflation model** |  |  |  |  |  |
| **Parameter** | **Estimate** | **z-value** | **CI<sub>low</sub>** | **CI<sub>up</sub>** | **p-value** |
| Intercept | -1.518 | -1.817 | -3.155 | 0.119 | 0.069 |
| Scat | -0.158 | -0.143 | -2.314 | 1.998 | 0.886 |
| Delta SVL | -0.662 | -1.490 | -1.533 | 0.209 | 0.136 |

**Table A3.** Parameter estimates and test statistics for the linear mixed model looking at deep breaths per second. The model included session and animal identity as random effects. CI – confidence interval.

| Parameter | Estimate | df | t-value | CI<sub>low</sub> | CI<sub>up</sub> | p-value |
| --- | --- | --- | --- | --- | --- | --- |
| Intercept | 0.489 | 35.2 | 0.352 | -2.142 | 3.112 | 0.727 |
| Scat | 0.015 | 315.8 | 0.559 | -0.036 | 0.065 | 0.576 |
| Water control | 0.023 | 315.3 | 0.727 | -0.038 | 0.085 | 0.468 |
| Own odour | -0.005 | 315.5 | -0.151 | -0.066 | 0.057 | 0.880 |
| Male | 0.108 | 18.9 | 1.879 | -0.004 | 0.219 | 0.076 |
| Stimulus order | -0.019 | 315.4 | -1.191 | -0.050 | 0.012 | 0.235 |
| Trial | -0.025 | 39.0 | -1.117 | -0.070 | 0.023 | 0.271 |
| Temperature | -0.006 | 35.1 | -0.118 | -0.108 | 0.096 | 0.907 |

**Table A4.**
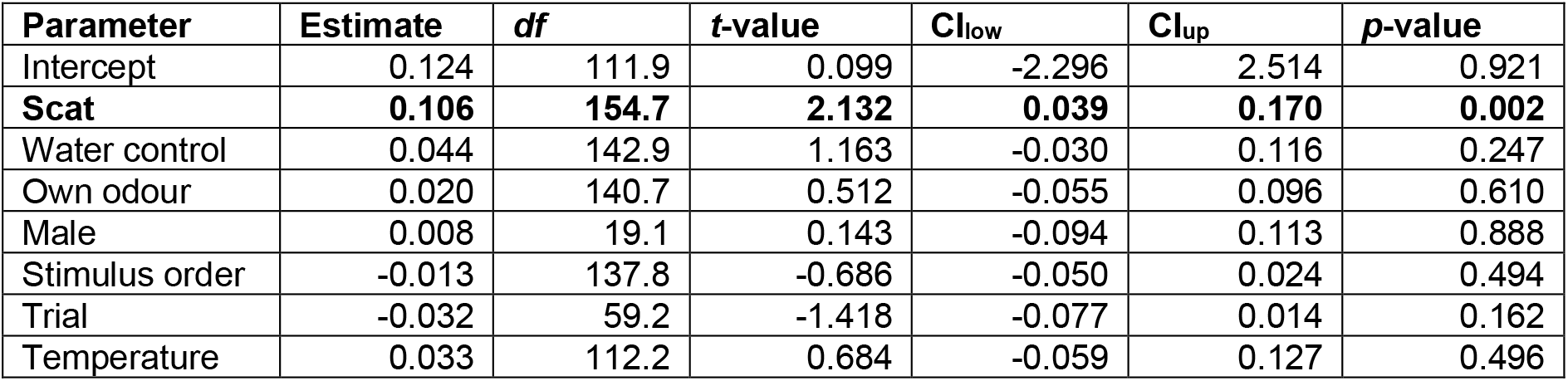
Parameter estimates and test statistics for the linear mixed model looking at breaths per second. The model included a random intercept of animal identity and a random slope of session. Significant results are highlighted in bold. CI – confidence interval.

**Table A5.**
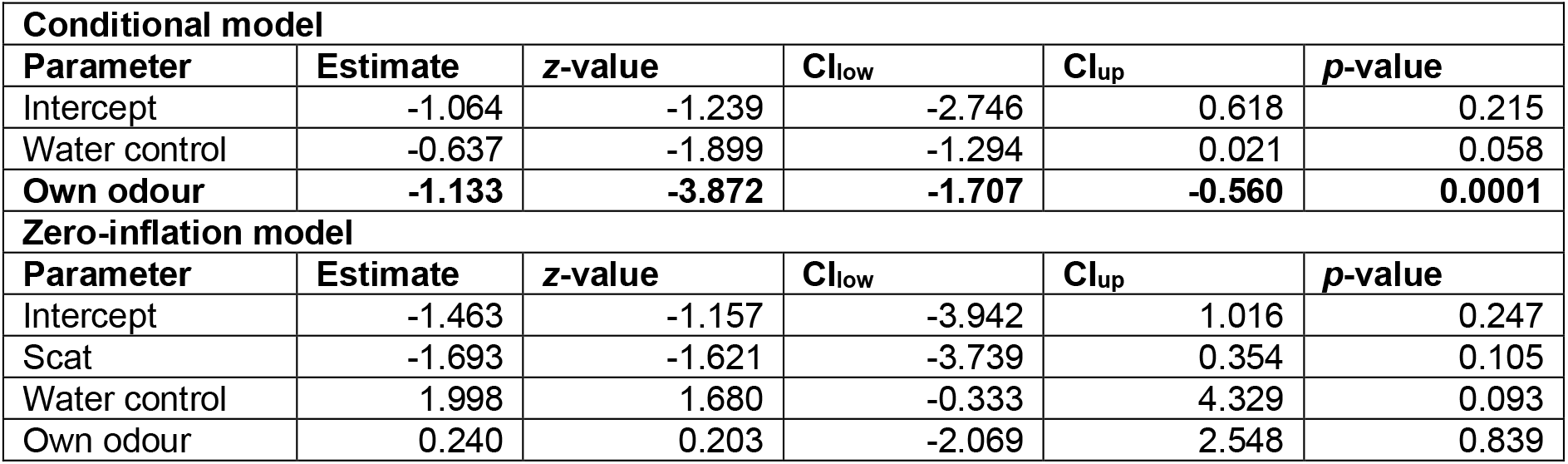
Parameter estimates and test statistics for the generalised linear mixed zero-inflation Poisson model looking at all ground tongue flicks across stimulus conditions in females. The conditional model included treatment, trial and animal identity as random effects, the zero-inflation model included animal identity as the random effect. Significant results are highlighted in bold. CI – confidence interval.

**Table A6.** Parameter estimates and test statistics for the generalised linear mixed zero-inflation Poisson model looking at all swab tongue flicks across stimulus conditions in females. The conditional model included treatment, trial and animal identity as random effects, the zero-inflation model included animal identity as the random effect. Significant results are highlighted in bold. CI – confidence interval.

| **Conditional model** |  |  |  |  |  |
| --- | --- | --- | --- | --- | --- |
| **Parameter** | **Estimate** | **z-value** | **CI<sub>low</sub>** | **CI<sub>up</sub>** | **p-value** |
| Intercept | -0.483 | -0.838 | -1.613 | 0.647 | 0.402 |
| **Water control** | **-1.405** | **-4.614** | **-2.002** | **-0.808** | **3.96\*10<sup>-6</sup>** |
| **Own odour** | **-0.927** | **-2.979** | **-1.538** | **-0.317** | **0.003** |
| **Zero-inflation model** |  |  |  |  |  |
| **Parameter** | **Estimate** | **z-value** | **CI<sub>low</sub>** | **CI<sub>up</sub>** | **p-value** |
| Intercept | -1.018 | -0.895 | -3.249 | 1.213 | 0.371 |
| Scat | 1.912 | 1.597 | -0.435 | 4.259 | 0.110 |
| Water control | -21.217 | -0.002 | -16347.2 | 16304.9 | 0.998 |
| Own odour | -1.809 | -1.445 | -4.262 | 0.645 | 0.148 |

